# Analysis of Suicidal Patients Admitted to The Emergency Rooms and Their Intensive Care Requirements, A Double-Center Study in Turkey

**DOI:** 10.1101/517417

**Authors:** Kocamer Şimşek Betul, Kocamer Şahin Şengül

**Author notes:** Betül Kocamer Şimşek (corresponding author), 05326105381. Postal Address: İncilipinar mah. Ali fuat cebesoy bulv. No.45 Şehitkamil Gaziantep Şengül Kocamer Şahin, 05336636876.

## Abstract

**Purpose:** In the present study, the clinical and socio-demographic data of the patients who admitted to the emergency department due to suicide attempt, the duration at the emergency department, and hospitalizations are examined. Requirement of intensive care and duration of hospitalization are investigated in the patients with suicide attempt.

**Methods:** Patients who were admitted to the emergency department of the hospitals after suicide attempts between 2015 and 2017 and earlier 2018 were included in the retrospective study. Reason for suicide, suicide modality, duration between the suicide attempt and arrival to the emergency department, suicide time, first treatment at the emergency department, hospitalization, mortality, and the levels at the intensive care unit (ICU) were retrospectively reviewed and analyzed.

**Results:** In the present study, 428 patients were included. Ratio of the female to male patients was 319/109. The mean age of the patients was 29.18±10.48. Most of the patients were single. The patients were mostly unemployed. Ninety-four (22.87%) patients were diagnosed with a psychiatric disorder. Four hundred twenty-two (98.59%) of the patients were attempted suicide with drugs/toxics. One hundred ninety-seven patients (49.75%) reported domestic violence and family issues reasons for suicide. Mean duration between the time of suicide and the time to arrive to the emergency department was 100.53±91.82 minutes. One hundred thirty (30.5%) patients were transferred to ICU, and 45 (10.5%) patients were followed in clinical departments. One hundred twenty (92.3%) patients hospitalized in the first-level ICU, 4 (3%) in the second-level ICU, and 6 (4.6%) in the third-level ICU. The mean ICU stay was 2.37±1.48 days.

**Conclusion:** The suicide attempts were prominent in acute poisoning cases. Majority of the patients stated domestic violence and family issues as a reason of suicide. They were discharged mostly from the emergency department and 10.5% of the patients were kept under surveillance in the departments. When the suicide attempts were evaluated in terms of their time, they were observed during day time at a higher rate.

## Introduction

Suicide is a public health problem that concerns communities, provinces and all the countries. More than 800.000 people die each year due to suicide. Suicide is the second most important cause of death between 15- and 29-years old patients worldwide (1). Methods of suicide attempt vary according to the countries. The most frequent suicide methods are overdosage, hanging themselves, asphyxiation, jumping out, pesticide poisoning, and use of firearms (2). Pesticide intake is one of the most common methods of suicide attempt globally (1) Most frequently used two methods of suicide leading to death in Turkey are hanging oneself and the use of firearms (3).

The patient with suicide attempt can be discharged or hospitalized based on the examination. The need for medical attention and interventions after a suicide attempt varies from 22% to 88% (4). Treatment settings include the department, intensive care unit, and the outpatient treatments. Severe suicide attempts, refusal of treatment, or mental state changes with metabolic, toxic, infectious situations or any other etiology generally require hospitalization (5). In Nordic countries, the rate of the patients requiring intensive care only after overdose varies between 3% and 15.7% (6). There is a limited number of studies in which suicidal attempts and the follow-up of the treatment at the intensive care unit are considered and there are no large-scale studies including the interventions at the emergency department and the follow-ups at the intensive care unit in Turkey.

In the present study, the clinical and socio-demographic data of the patients with suicide attempt who admitted to the emergency department, duration of admission to the emergency department, and hospitalizations are examined. Requirement of intensive care and duration of hospitalization are investigated in the patients with suicide attempt.

## Materials and Methods

### Data collection

The present study was retrospectively conducted at Adana State Hospital and Sanko University Hospital of Faculty of Medical. Approval for the study was granted by the Institutional Ethics Committee of Sanko University, Gaziantep, Turkey (No: 2017/01-4 date:25.01.2017).

Patients who were admitted to emergency departments of the hospitals after suicide attempts between 2015 and 2017 and earlier 2018 were included in the study. The names, phone numbers and epicrisis of the patients were retrospectively screened from the archives of the hospitals.

Age, gender, relationship status, employment status, alcohol or substance abuse, history of psychiatric disorders, history of chronic illnesses, reason for suicide, suicide modality, previous suicide attempts, history of suicide in family members, duration between the suicide attempt and arrival to the emergency department, suicide time, initial treatment at the emergency department, hospitalization, mortality, and intensive care unit (ICU) levels were retrospectively reviewed and analyzed. If the answers for the said elements were not present in the hospital registries, the patients were called, and the missing questions were asked. Some of the patients did not want to talk or some of them could not be reached, and therefore, they were recorded as missing data.

### Statistical analysis

SPSS 25.0 (IBM Corporation, Armonk, New York, United States) program was used to analyze the variables. Chi-Square and Binominal tests are used for the homogeneity of categorical variables, Fisher-Freeman-Holton tests are used for the comparison with each other and they are tested by using the Monte Carlo Simulation technique. The ratios of the columns are compared with each other and expressed according to the Bonferroni corrected p value results. Quantitative variables were shown as mean ± SD (Standard Deviation) and median (Minimum/maximum) and categorical variables as n (%) in the tables. The variables were examined at confidence level of 95% and p value less than 0.05 was accepted as significant.

### Results

In the present study, 428 patients were included. Ratio of the female to male patients was 319/109. The mean age of the patients was 29.18±10.48. Most of the patients were married (n=189) and single (n=205) and others were widow (n=20) or divorced (n=14). The patients were mostly unemployed. One hundred nine patients were housewives (n=109), 103 were working in a job, 58 were students and 3 were retired.

Table 1 shows the sociodemographic data of the patients.

**Table 1.** Demographic data. *p<0.001

|  | n (%) |  |  |
| --- | --- | --- | --- |
| Gender |  |  |  |
| Female | 319 (74.53)* |  |  |
| Male | 109 (25.47) |  |  |
| Relationship status |  |  |  |
| Single | 205 (47.90)* |  |  |
| Married | 189 (44.16)* |  |  |
| Widow | 20 (4.67) |  |  |
| Divorced | 14 (3.27) |  |  |
| Occupying |  |  |  |
| Unemployed | 136 (33.25)* |  |  |
| Working in a job | 103 (25.18) |  |  |
| Housewife | 109 (26.65) |  |  |
| Student | 58 (14.18) |  |  |
| Retired | 3 (0.73) |  |  |
|  | N | Mean±SD. | Median (Min. / Max.) |
| Age | 428 | 29.18±10.48 | 26 (14 / 67) |

Ninety-four patients were diagnosed with a psychiatric disorder, and ninety of them were receiving treatment. Sixteen patients were diagnosed with chronic illnesses as diabetes mellitus, hypertension, cardiac problems and cancer, and 3 of these patients were diagnosed with neurological disorders. Fifteen patients had substance abuse and 31 patients had alcohol abuse. Eighty-five patients had attempted suicide previously, 23 of these patients had a history of suicide attempt in the family members and 21 of them had a history of psychiatric disorders in the family members. Medical history of the patients was analyzed, and they are shown in table 2.

**Table 2.** Medical history of patients. *p<0.001 **Other comorbidities are diabetes mellitus, hypertension, cardiac problems and cancers.

| History | n (%) |
| --- | --- |
| Alcohol abuse |  |
| Yes | 31 (7.29) |
| No | 394 (92.71)* |
| Substance abuse |  |
| No | 408 (96.45)* |
| Yes | 15 (3.55) |
| Comorbidities |  |
| None | 409 (96.23)* |
| Other ** | 13 (3.06) |
| Neurological | 3 (0.71) |
| Psychiatric disorder |  |
| None | 317 (77.13)* |
| Yes | 94 (22.87) |
| Psychiatric treatment |  |
| None | 293 (76.50)* |
| Yes | 90 (23.50) |
| History of suicide attempt before |  |
| No | 335 (79.76)* |
| Yes | 85 (20.24) |
| History of suicide attempt in family members |  |
| No | 353 (93.88)* |
| Yes | 23 (6.12) |
| Psychiatric disorder in family members |  |
| No | 341 (94.20)* |
| Yes | 21 (5.80) |

Four hundred twenty-two of the patients were attempted suicide with drugs/toxic substances. Of these patients, 3 attempted suicide with cutting tools, 2 by jumping over (1=4^th^ floor, 1=2^nd^ floor), and 1 by blowing his/her brain out (died within 12 hours). The reasons for attempting suicide were analyzed, and 396 patients specified the reason, but 32 patients did not want to give information. In the said 396 patients, majority stated the reason as domestic violence and family issues (n=197) followed by chronic illnesses, (n=64) and others specified the reason as communication problems (n=46), sexual problems (n=28), economic problems (n=18), loneliness (n=12), alcohol/substance abuse (n=9), parental conflicts (n=8), exams (n=8), death/lost (n=6), juvenile problems (3), and school (n=2). Table 3 shows the suicide modality of the patients.

**Table 3.** suicide descriptives. *p<0.001

|  |  |
| --- | --- |
| **Suicide modality** |  |
| Drugs/toxics | 422 (98.59)* |
| Cutting tools | 3 (0.70) |
| High jump | 2 (0.47) |
| Gun | 1 (0.24) |
| **Reason for suicide attemp** |  |
| Domestic violence and family issues | 197 (49.75)* |
| Chronic illnesses | 64 (16.16) |
| Communication problems | 46 (11.62) |
| Sexual problems | 23 (5.81) |
| Economical problems | 18 (4.54) |
| Loneliness | 12 (3.03) |
| Alcohol/substance abuse | 9 (2.27) |
| Parental conflict | 8 (2.02) |
| Exams | 8 (2.02) |
| Death/lost | 6 (1.52) |
| Juvenile problems | 3 (0.76) |
| School | 2 (0.51) |
| **Suicide time** |  |
| [24:00-06:00] | 75 (18.9)* |
| [06:00-18:00] | 174 (43.9) |
| [18:00-24:00] | 147 (37.1) |

The times of suicide attempts were examined in three different periods: evening period (18:00-23:59), night period (00:00-05:59), and day time period (06:00-17:59). The records of the times of suicide attempts were available in 396 patients. Most of the patients attempted suicide during the day time period (n=174), secondly in the evening period (n=147), and finally in the night period (n=75) (table 3).

Table 4 shows the descriptive characteristics of ICU and the emergency department. Four hundred twenty patients were treated with gastric lavage and activated charcoal in the emergency department, 3 patients with tracheal intubation and transferred to ICU, and 3 patients with tracheal intubation and then immediately transferred to the operating room. One hundred twenty (92.3%) patients hospitalized in first-level ICU, 4 (3%) in second-level ICU, and 6 (4.6%) in third-level ICU.

**Table 4.** ICU and emergency room descriptives. *p<0.005

|  |  |  |
| --- | --- | --- |
| **Steps in ICU** |  | n (%) |
|  | 1.Step | 120 (92.3) * |
|  | 2. Step | 4 (3) |
|  | 3. Step | 6 (4.6) |
| **Stay in hospital** |  |  |
|  | Discharge from emergency room | 251 (58.9)* |
|  | ICU | 130 (30.5) |
|  | Clinical Services | 45 (10.5) |
| **Treatment in emergency room** |  |  |
|  | Gastric lavage and activated charcoal | 420 (98.5)* |
|  | Tracheal intubation and transfer to ICU | 3 (0.7) |
|  | Tracheal intubation and transfer to operating room | 3 (0.7) |
| **Treatment in ICU** |  |  |
|  | Monitored Care | 120 (92.3) * |
|  | Hemodialysis | 4 (3) |
|  | Mechanical ventilation and Hemodialysis | 3 (2.3) |
|  | Mechanical ventilation and operation | 3 (2.3) |
| **Discharge from ICU** |  |  |
|  | Discharged | 129 (99.24)* |
|  | Death | 1 (0.76) |

Two hundred fifty one patients were discharged from the emergency department, 130 patients were transferred to ICU, and 45 patients were followed in clinical departments. Mean duration between the time of suicide attempt and the time to arrive to the emergency department was 100.53±91.82 minutes (n=416) (table 5).

**Table 5.** Time to reach to the emergency room and Stay in ICU.

|  | **Mean±SD.** | **Median (Min. / Max.)** |
| --- | --- | --- |
| **Time to reach to the emergency room (min)** | 100.53±91.82 | 60 (6 / 540) |
| **Stay in ICU (day)** | 2.37±1.48 | 2 (1 / 12) |

Sixty-three female and 40 male patients were working in a job, and 92 female patients and 44 male patients were unemployed (table 6).

**Table 6.** Relationship between gender and occupying

|  | **Female**<br>**n** | **Male**<br>**n** |
| --- | --- | --- |
| **Working in a job** | 63 | 40 |
| **Unemployed** | 92 | 44 |
| **Retired** | 1 | 2 |
| **Housewife** | 109 | 0 |
| **Student** | 41 | 17 |

The reasons for suicide attempts were mostly domestic violence and family issues (52.7%) in female patients and were mostly the economic problems (13.5%) and illnesses (13.5%) in male patients. The most common cause in all relationship status when the relationship status and the reason of suicide are compared is domestic violence and domestic problems. Similarly, the most common causes in employment conditions when the employment status and the reason of suicide are compared, are domestic violence and domestic problems. Table 7 shows the relationship status, and the occupational status.

**Table 7.** Relationship between suicide reasons and gender, relationship status, occupying *p<0.001

|  | Gender |  | Relationship status |  |  |  | Occupying |  |  |  |  |
| --- | --- | --- | --- | --- | --- | --- | --- | --- | --- | --- | --- |
|  | Female<br>n (%) | Male<br>n (%) | Single<br>n (%) | Divorced<br>n (%) | Widow<br>n (%) | Married<br>n (%) | Working<br>in a job<br>n (%) | Unemployed<br>n (%) | Retired<br>n (%) | House<br>wife<br>n (%) | Student<br>n (%) |
| Domestic violence and family issues | 158 (52.7)* | 39 (40.6) | 69 (37.3)* | 9 (64.3)* | 7 (41.2)* | 112 (62.2)* | 51 (52) | 51 (40.8) | 1 (50) | 64 (61)* | 20 (39.2) |
| Alcohol/substance abuse | 1 (0.3) | 8 (8.3) | 9 (4.9) | 0 (0) | 0 (0) | 0 (0) | 3 (3.1) | 4 (3.2) | 0 (0) | 0 (0) | 2 (3.9) |
| Sexual problems | 15 (5) | 8 (8.3) | 16 (8.6) | 0 (0) | 1 (5.9) | 6 (3.3) | 5 (5.1) | 6 (4.8) | 0 (0) | 5 (4.8) | 5 (9.8) |
| Parental conflict | 7 (2.3) | 1 (1) | 7 (3.8) | 0 (0) | 0 (0) | 1 (0.6) | 0 (0) | 7 (5.6) | 0 (0) | 0 (0) | 1 (2) |
| Economic problems | 5 (1.7) | 13 (13.5)* | 7 (3.8) | 0 (0) | 0 (0) | 11 (6.1) | 6 (6.1) | 9 (7.2) | 0 (0) | 1 (1) | 1 (2) |
| Juvenile problems | 2 (0.7) | 1 (1) | 3 (1.6) | 0 (0) | 0 (0) | 0 (0) | 0 (0) | 0 (0) | 0 (0) | 0 (0) | 3 (5.9) |
| Chronic illnesses | 51 (17) | 13 (13.5)* | 29 (15.7) | 3 (21.4) | 5 (29.4) | 27 (15) | 22 (22.4) | 19 (15.2) | 1 (50) | 18 (17.1) | 4 (7.8) |
| Communication | 35 (11.7) | 11 (11.5) | 26 (14.1) | 1 (7.1) | 1 (5.9) | 18 (10) | 11 (11.2) | 15 (12) | 0 (0) | 12 (11.4) | 7 (13.7) |

| problems |  |  |  |  |  |  |  |  |  |  |  |
| --- | --- | --- | --- | --- | --- | --- | --- | --- | --- | --- | --- |
| School | 2 (0.7) | 0 (0) | 1 (0.5) | 0 (0) | 0 (0) | 1 (0.6) | 0 (0) | 0 (0) | 0 (0) | 0 (0) | 2 (3.9) |
| Death/lost | 6 (2) | 0 (0) | 2 (1.1) | 0 (0) | 2 (11.8) | 2 (1.1) | 0 (0) | 2 (1.6) | 0 (0) | 4 (3.8) | 0 (0) |
| Exams | 6 (2) | 2 (2.1) | 8 (4.3) | 0 (0) | 0 (0) | 0 (0) | 0 (0) | 5 (4) | 0 (0) | 0 (0) | 3 (5.9) |
| Loneliness | 12 (4) | 0 (0) | 8 (4.3) | 1 (7.1) | 1 (5.9) | 2 (1.1) | 0 (0) | 7 (5.6) | 0 (0) | 1 (1) | 3 (5.9) |

## Discussion

It is critical to understand the extent of suicide attempts in order to develop the protection and intervention programs. The present study contributes to the literature regarding the suicide modalities, duration between the suicide attempt and arrival to the emergency department, time of the suicide attempt, first treatment at the emergency department, hospitalization, and the intensive care unit (ICU) stages in order to monitor suicide-related treatment level.

Previous studies found that suicide attempts are higher in females and males are more likely to have completed suicide (7,8). Similarly, in the present study, the number of suicide attempts were higher in women than in men and 1 person who died due to a suicide attempt was male. Also, being single acts as a risk factor for a suicide attempt (9). Consistently, it was found that suicide attempts were higher in singles.

In the present study, the predominant method for suicide attempt was drug intoxication, and the second reason was slitting his/her wrists. Only 3 patients attempted suicide by slitting their wrists. This frequency is compatible with the previous studies regarding suicide attempt (8).

Psychiatric disorders, history of suicidal behaviors, and substance abuse are important risk factors for suicide attempt (10). In the current study, 94 patients were diagnosed with a psychiatric disorder, 15 had substance abuse, 31 had alcohol abuse, and 85 had a previous suicide attempt. The approaches to interpersonal/relationship problems, psychiatric morbidity may be complementary for the doctors at the emergency department as well as the psychiatrists.

There is a significant correlation between domestic violence and suicide attempt tendency in developing countries (11). In the current study including 396 patients, most of the patient stated domestic violence and family issues (n=197) as the reason for a suicide attempt followed by chronic illnesses.

The results of the studies regarding the times of suicide attempt are not consistent. In the present study, the highest incidence rate for suicide attempt was the day time period (06:00-17:59). Times of suicide attempts are analyzed in different time intervals (ex. Most frequent time intervals according to the studies are 15.00-18.00, 06.00-16.00, 12.00-16.00, 8.30-12.30) (12-14), it may be considered that the suicide attempts are more frequent when the patient is awake in the day time.

One of the distinctive results of the present study is how long the patients reach the emergency department after suicide attempt. In a study conducted in Turkey, the median time from the exposure to substance intake to the ED was 2 hours (17). It took 100.53±91.82 minutes to arrive to the emergency department after any suicide attempt in the current study. 130 (30.5%) patients were transferred to ICU, and 45 (10.5%) patients were followed in the clinical departments. The mean ICU stay was 2.37±1.48 days.

The patients who needed intensive care due to suicide attempts were also examined. In a previous study, it was determined that 12.8% of the patients, who applied due to suicide attempt, were admitted to ICU (16). In the present study, 130 (30.5%) patients were transferred to ICU. According to the classification of the ICU levels, 120 patients with suicide attempts, who took drugs, were kept in ICU at first level, 4 patients at the second level (one needed CPAP ventilation, two needed hemodiafiltration, 1 had a Glasgow coma scale of 10). Mechanical ventilation was required for 6 patients, so they were accepted as the ICU level 3. Two of these patients jumped from a high place, 1 attempted suicide with a gun, and 3 had a respiratory failure due to the use of drugs. All the patients in the 1^st^ level were accepted to ICU due to the possible adverse events like arrythmia, loss of consciousness and respiratory, cardiac, renal or hepatic failures and they were monitored according to the said adverse events as mentioned in the previous studies (6).

In a study in Turkey, it was determined that the rate of deaths caused by the suicide attempts by using drugs is 6%. In another study, this rate was found to be 0.1% (17,18). According to the findings of World Health Organization, mortality rate after a suicide attempt is 11,4 per 100.000 (19). In the current study, it was found that 1 patient died by blowing his/her brains out in a suicide attempt. No patient died due to the suicide attempts by using drugs. The result supports the low mortality rate in the suicide attempts in Turkey.

## Conclusion

The suicide attempts were prominent in acute poisoning. Majority of the patients stated domestic violence and family issues as a reason of suicide. They were discharged mostly from the emergency department and 10.5% of the patients were kept under surveillance in the departments. When the suicide attempts were evaluated in terms of their time, they were observed during day time at a higher rate.

## Limitation

A major limitation of this study is the retrospective design. There are no detailed psychiatric examination findings in the emergency departments.

